# Modulation of beta oscillatory dynamics in motor and frontal areas during physical fatigue

**DOI:** 10.1101/2024.06.11.598466

**Authors:** Pierre-Marie Matta, Robin Baurès, Julien Duclay, Andrea Alamia

**Author notes:** **Corresponding author:** Pierre-Marie Matta.

## Abstract

Beta-band oscillations have been suggested to promote the maintenance of the current motor (or cognitive) set, thus signaling the ‘status quo’ of the system. While this hypothesis has been reliably demonstrated in many studies, it fails to explain changes in beta-band activity due to the accumulation of physical fatigue. In the current study, we aimed to reconcile the functional role of beta oscillations during physical fatigue within the status quo theory. Using an innovative EEG design, we identified two distinct beta-band power dynamics in the motor areas as fatigue rises: (i) an enhancement at rest, supposedly promoting the resting state, and (ii) a decrease during contraction, thought to reflect the increase in motor cortex activation necessary to cope with the muscular fatigue. We then conducted effective connectivity analyses, which revealed that the modulations during contractions were driven by frontal areas. Finally, we implement a biologically plausible model to replicate and characterize our results mechanistically. Together, our findings anchor the physical fatigue paradigm within the status quo theory, thus shedding light on the functional role of beta oscillations in physical fatigue. We further discuss a unified interpretation that might explain the conflicting evidence previously encountered in the physical fatigue literature.

## I. Introduction

Beta-band oscillations (∼13-30Hz) have been associated with a wide range of cognitive and sensorimotor processes (Barone & Rossiter, 2021; Lundqvist et al., 2024). Regarding the latter, beta oscillations prevail particularly in the absence of movements, whereas performing a contraction causes a drop in their power (Kilavik et al., 2013). In line with these findings, some authors have suggested that beta oscillations reflect an ‘idling’ state of the system (Pfurtscheller et al., 1996), a ‘status quo’ that promotes the maintenance of the current motor (or cognitive) state (Engel & Fries, 2010). Accordingly, a motor activation can be interpreted as a change in the system (Formaggio et al., 2008; Stevenson et al., 2011; Yuan et al., 2010), which induces a decrease in beta power.

While these theories proved reliable explanatory power in several studies, they fail to successfully explain experimental conditions in which beta oscillations play a crucial role (Schmidt et al., 2019), such as physical fatigue (i.e., decreased ability to exert muscle force (Gandevia, 2001)). Specifically, previous work investigating the modulation of beta oscillations during submaximal exercises revealed an increase in beta oscillatory power in the motor cortex (MC) following a rise in fatigue levels (Bailey et al., 2008; Ciria et al., 2018; Enders et al., 2016; Kubitz & Mott, 1996; Lin et al., 2021; Suviseshamuthu et al., 2021; Yang et al., 2009). This gain in beta-band power has been interpreted as an increase in MC activation (Enders et al., 2016; Kubitz & Mott, 1996; Lin et al., 2021; Yang et al., 2009), necessary to recruit additional motor units to overcome the growing muscle fatigue (Enoka & Stuart, 1992) when sustaining a submaximal exercise (Benwell et al., 2007; Liu et al., 2003; Taylor et al., 2016). However, these experimental results are at odds with the status quo theory, which would predict a decrease in beta-band oscillatory power, due to an increase (i.e., a change) in the MC activity necessary to pursue the physical effort.

In the current study, using an experimental design optimized for our purposes, we address such discrepancy and provide an interpretation that reconciles conflicting experimental evidence with our current understanding of beta oscillations during fatigue and the status quo theory. Specifically, we recorded electroencephalography (EEG) signals to investigate how physical fatigue modulates the beta oscillatory dynamics in the MC and the prefrontal cortex (PFC). Participants performed 100 isometric knee extensions against a fixed submaximal resistance in two distinct temporal conditions (either 10 s or 12 s contractions), aiming to generate two different levels of fatigue (Fig 1A). Importantly, this design allowed us to study the evolution of fatigue neural correlates with a relatively high temporal resolution while minimizing fatigue-related muscle artifacts and confounds such as muscular compensation. In addition, the relatively low submaximal load chosen (≤ 20% of maximal force) and the endurance of the muscle engaged (quadriceps) will enable us to observe the early stages of fatigue, where we will be able to dissociate beta-band dynamics during the contraction and rest. These two temporal moments will allow us to observe different dynamics. On the one hand, in line with previous studies and the status quo theory, we hypothesized that the beta-band power would increase with physical fatigue during rest (i.e., between each contraction), reflecting a growing promotion of the resting state. On the other hand, different than in earlier studies but consistent with the status quo theory, we predicted that the beta-band power would decrease over the motor areas during the contraction with physical fatigue, highlighting the increasing MC activation required to cope with the muscle fatigue.

**Figure 1.**
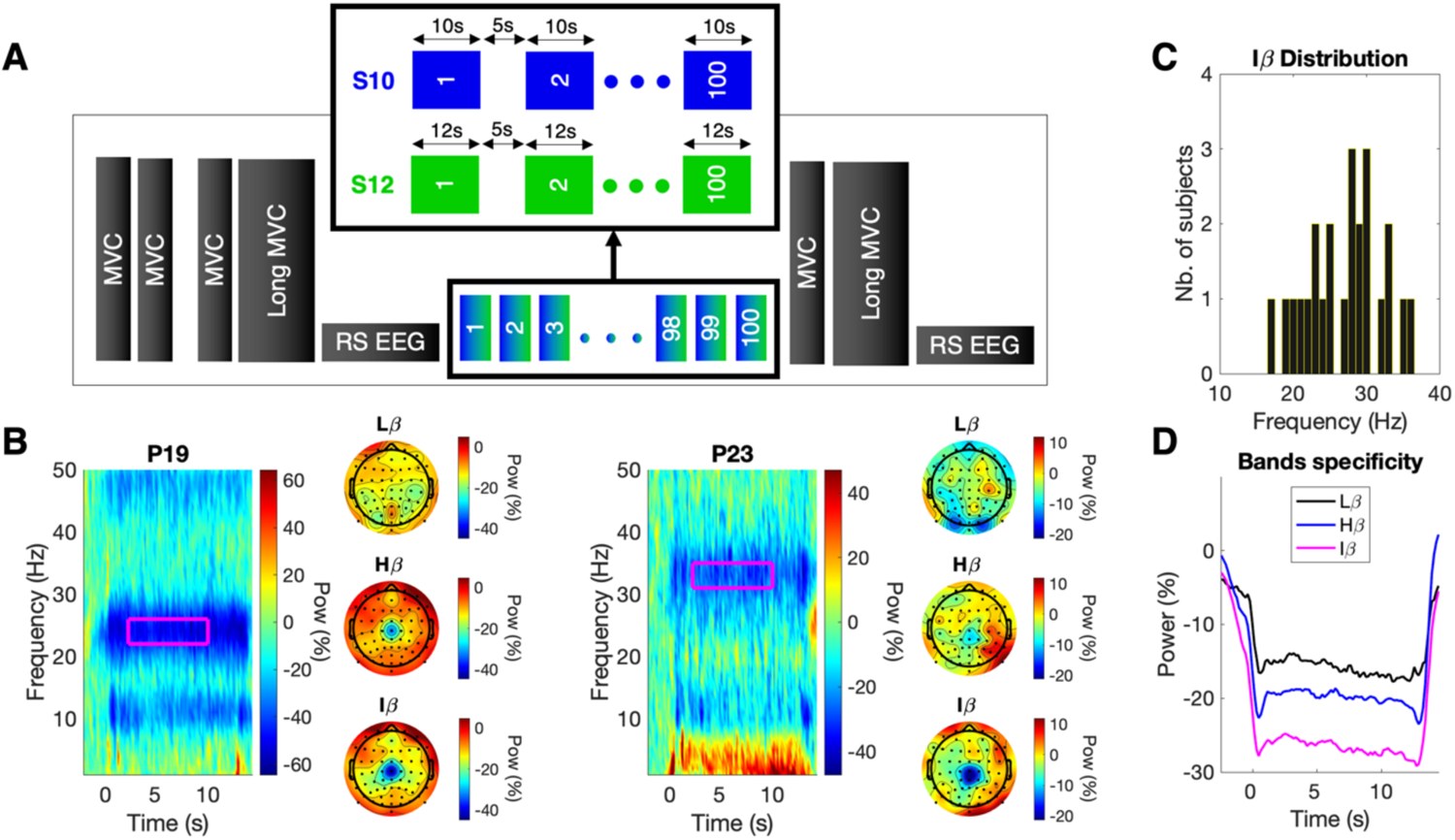
Experimental protocol and individualized beta-band computation. A. Procedure. All participants performed 100 contractions during two separate sessions (S). Each contraction lasted 10 or 12 s, depending on the session (S10 and S12, respectively), and was separated by a fixed rest time (5 s). EMG and EEG activity were recorded during the 100 contractions. Before and after the task, Maximal Voluntary Contractions (MVC, ∼3 s) and long MVC (∼30 s) were performed to assess the fatigue generated by the 100 contractions. Resting State (RS) EEG (3 min) was recorded before and after the task to quantify the associated EEG correlates. B. Spectrogram representations of Cz electrode for two typical participants (P) in the S12 session. Each plot represents the median over 100 contractions after a baseline correction between the -3 to -1 s period preceding the contraction (contraction onset = 0 s). Magenta rectangles represent the individualized beta band, which ranges from -2 to +2 Hz around the frequency with the largest desynchronization in the period of interest (2 to 10 s) for each participant. Note that Iβ was computed accounting for both S10 and S12 sessions (see Methods for details). Only the S12 session is represented here for visual purposes. Topographic representations next to each spectrogram have been obtained averaging over the period of interest (2 to 10s) for Low-Beta (Lβ), High-Beta (Hβ), and Individualized-Beta (Iβ). C. Iβ distribution over frequencies. The Iβ extraction procedure leads to optimal individual desynchronization frequencies ranging from 17 to 36 Hz among participants. D. Beta-band desynchronization. Normalized spectral power from the Cz electrode during the S12 session was extracted for Lβ, Hβ, and Iβ and averaged over contractions and participants. We can observe (i) a larger desynchronization in Hβ compared to Lβ, reflecting Hβ specificity to lower limb contraction, (ii) a greater desynchronization in Iβ compared to Hβ, highlighting the importance of extracting a narrower individualized beta-band to avoid accounting for frequencies non-involved in the desynchronization. We applied a moving average window for visualization purposes only.

## II. Results

Participants performed two sessions (spaced 3 to 10 days apart) of 100 quadriceps isometric contractions (10 or 12 seconds each) separated by a fixed rest time (5 s) (Fig 1A). The contraction intensity was identical in both sessions, corresponding to 20% (up to a fixed value, see Methods for details) of the participant’s maximal force. EEG and EMG were recorded throughout the task and analyzed specifically over a 2 to 10 s window within each contraction (0 s being the contraction onset) in both sessions. EEG was also considered during the task at rest, between subsequent contractions (from -3 to -1 s before contraction onset). Before and after the task, participants performed short (∼3 s) and long (30 s) Maximal Voluntary Contractions (MVC), as well as 3 minutes EEG resting states.

### Neuromuscular markers of physical fatigue

At first, we confirmed and quantified the physical fatigue generated by the task by comparing the maximal force produced by participants before and after the 100 contractions (pre- and post-task). As physical fatigue can manifest in both the maximal force production and the ability to sustain it, we implemented various measurements to determine which best captures its effect. Specifically, we looked at the pre-post changes in maximal force production capacity (‘Short Max’) and at the pre-post changes in the difference between the start and end of a long MVC (‘Long Max Diff’), as suggested in the literature (Lebesque et al., 2022). Additionally, concerning the long MVC, we derived the force integral of the long MVC over time (‘Long Max Int.’), which could provide a more accurate sustaining ability measure by accounting for the full range of effort. We verified which measures were the most sensitive to physical fatigue by testing their variation rate ((post-pre)/pre*100) against 0. Short Max and Long Max Int. were significantly below 0 in both sessions (p < .001, d > 1.86, BF > 1.07×10^+6^, for all, unilateral one-sample t-test, H1: DV < 0), indicating that the task generated a significant decline in the capacity to produce and maintain a maximal force (Fig 2A). We then used these two variables to assess whether the two sessions had a fatigue difference. Short Max only showed a significant difference between S10 and S12 (t_(23)_ = 1.84, p = .039, d = 0.38, BF = 1.75, unilateral pairwise t-tests, H1: S10 > S12), thus revealing a larger decline in the ability to produce a maximal force (i.e., more fatigue) after the 12 s contractions.

After quantifying the difference between pre- and post-task measures, we ran an RM-ANOVA on the amplitude of the electromyography (EMG) signals recorded during the 100 contractions to assess neuromuscular fatigue accumulation (Kim et al., 2007; Matta et al., 2024; Merletti et al., 1991). To this end, we considered the normalized averaged root-mean-square (RMS) of each surface quadriceps muscle, with BIN (from 1 to 10, each comprising ten contractions) and SESSION (S10, S12) as fixed factors. We found a robust BIN effect (F_(9,207)_ = 41.49, p < .001, η^2^ = 0.64, BF = 3.30×10^+38^), indicating increasing fatigue over contractions, as well as a substantial BIN-SESSION interaction (F_(9,207)_ = 3.43, p < .001, η^2^ = 0.13, BF = 28.43), highlighting a growing fatigue difference between the two sessions (with S12 > S10) (Fig 2B).

### Beta-band modulations in motor areas during the task

Once we confirmed through the force and EMG signals that physical fatigue increased over contractions, we investigated MC beta-band power dynamics. In line with the literature, we considered Cz to reflect the MC activity associated with the quadriceps contraction (Dal Maso et al., 2012; Forrester et al., 2006). Spectral power was quantified in an Individualized-Beta (Iβ) band, defined as the frequency presenting each participant’s largest desynchronization during contraction (Fig 1B). Iβ was extracted during contraction and at rest (i.e., between contractions), and normalized by the average power of the first contractions (see Methods for details).

To investigate the motor-related neural modulations during the contractions, we first ran RM-ANOVAs with BIN (1 to 10) and SESSION (S10, S12) as fixed factors on Iβ Cz. Our results revealed a BIN effect for Iβ Cz (F_(9,207)_ = 2.67, p = .006, η^2^ = 0.10, BF = 5.55), highlighting a decreasing power over the task, while no SESSION or BIN-SESSION interaction effects were found (p > .27, BF < .33, for both factors) (Fig 2B). We surmised that this beta-band power decrease reflects the progressive increase in the MC activation (Pfurtscheller & Lopes da Silva, 1999) to cope with the rising muscular fatigue. Additionally, we also performed the same analysis on Iβ Cz quantified during the period of rest between each contraction to account for fatigue-related changes in the beta-band power resting baseline. Importantly, we also found a BIN effect for Iβ Cz recorded at rest (F_(9,207)_ = 3.04, p = .002, η^2^ = 0.12, BF = 2.71), showing an increase in beta power over the task (Fig 2B). Remarkably, this result reveals an opposite pattern to what we observed during the contractions, suggesting two distinct dynamics for beta-band oscillations during fatigue: in line with the status quo theory, these results highlight, on the one hand, an increasing power at rest, which promotes the resting state, and, on the other hand, a decreasing power during contractions reflecting the growing MC activation. Finally, there were no SESSION or BIN-SESSION interaction effects regarding Iβ Cz at rest (p > .51, BF < .45, for both).

**Figure 2.**
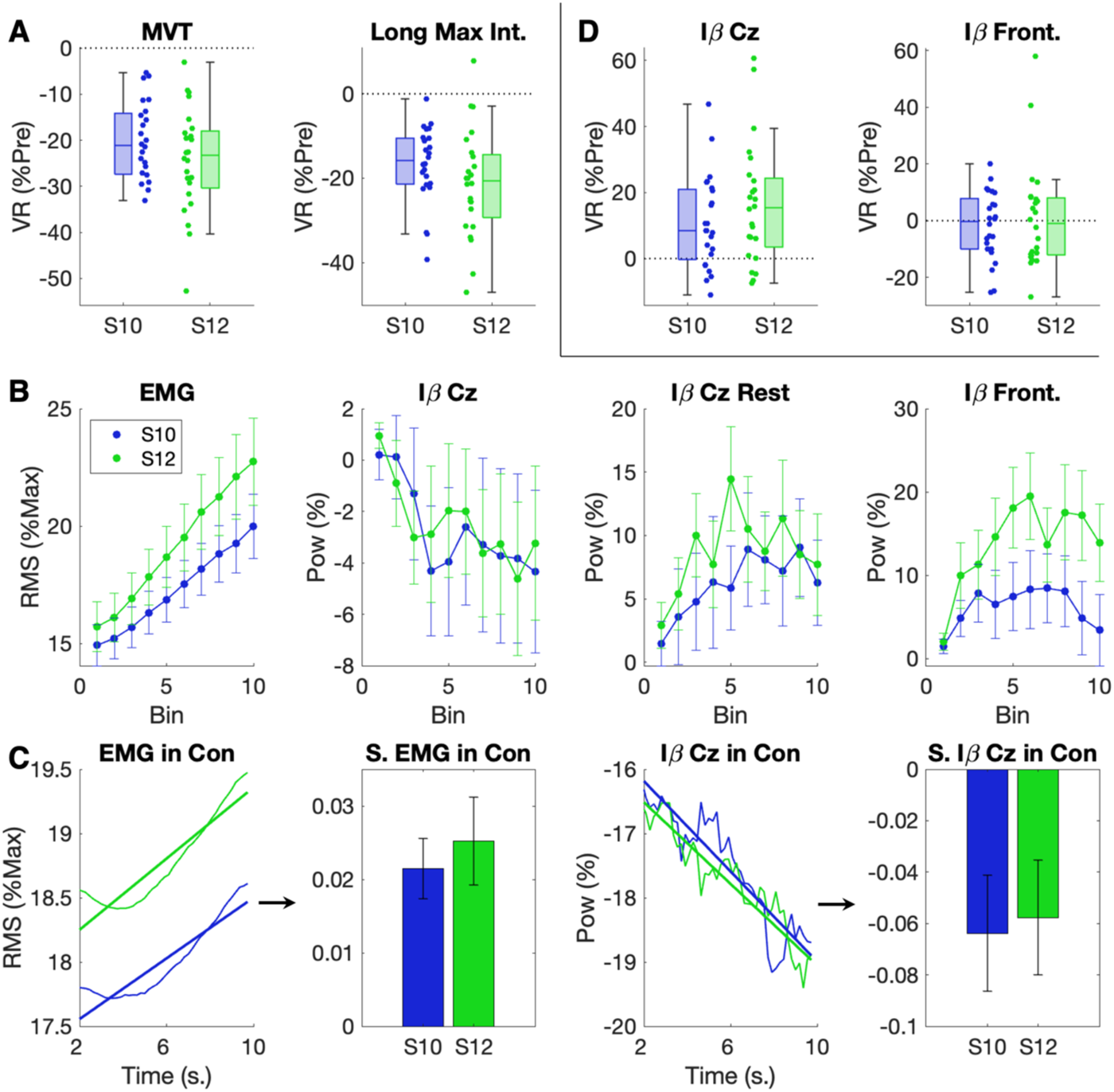
Neuromuscular fatigue assessment and beta-band power modulations. A. Neuromuscular Pre-Post Variation Rates (VR). A decrease in Maximal Voluntary Torque (MVT) reflects a decline in the maximal force production, while a reduction in Long Max Integral reveals a decline in the ability to sustain a maximal force over a given duration (30 s here). Both measurements allow for the assessment of physical fatigue. B. EMG and EEG modulations over the main task. Panel 1 represents the EMG expressed in normalized RMS over contractions, whose increase reflects neuromuscular fatigue accumulation. Panel 2 and 3 represent relative power changes (respective to the first contractions, see Methods for details) quantified in an Individualized-Beta band (Iβ) in Cz, in contraction (panel 2), and at rest (panel 3), that is, between contractions. Panel 4 exhibits the power-relative changes in frontal electrodes quantified during contractions. The dots represent the averaged distribution for all panels, and the error bars are the standard error mean (SEM). C. EMG and Iβ Cz during contractions. The lines represent the average over participants of EMG and Iβ Cz during a contraction, with the straight lines representing their slopes. For representation only, signals were smoothed with a moving average window. The bar plots display the average slope extracted for each participant and session, and the error bars are the SEM. D. EEG Pre-Post Variation Rates. Iβ changes in Post relative to Pre were quantified in Cz and frontal electrodes.

### Neural motor-related changes within contraction

We then questioned whether there was a fatigue effect within each contraction. Specifically, we investigated the EMG signals and the motor-related beta-band modulations irrespective of the contraction number (i.e., averaging over the BIN factor). Accordingly, we averaged the 100 contractions (i.e., RMS for EMG and Iβ Cz for EEG) for each participant, and we computed the slope of EMG and Iβ Cz over our window of interest (from 2 to 10 s) to identify if their trend over time was significantly different from 0. We found a significant positive slope for EMG (p < .001, d > 0.86, BF > 91.65, for both sessions) and a negative one for Iβ Cz (p < .017, d > 0.53, BF > 3.14, for both sessions) (Fig 2C), hence mirroring the results obtained in the previous analyses over contractions. These results again support the status quo interpretation, as the increasing fatigue, reflected by the positive EMG trend, is associated with an enhanced beta-band desynchronization, thought to reflect the growing MC activation compensating for muscle fatigue.

### Beta-band power modulations in frontal areas during the task

We hypothesized that the beta-band power modulations revealed in motor areas could be caused by a top-down modulation from frontal areas. Indeed, the PFC shares extensive connections with the motor areas (Miller & Cohen, 2001) and plays a key role in physical effort maintenance (Bigliassi & Filho, 2022; Denis, 2020; McMorris et al., 2018; Robertson & Marino, 2016). To investigate the modulations occurring in the PFC, we considered together the normalized Iβ power of lateral-frontal (F7-F8) and mid-frontal (F3-F4) electrodes (Okamoto et al., 2004; Robertson & Marino, 2015) during each contraction. We performed an RM-ANOVA with BIN (1 to 10) and SESSION (S10, S12) as fixed factors on Iβ in frontal electrodes. We found a significant BIN effect (F_(9,207)_ = 3.57, p < .001, η^2^ = 0.13, BF = 31.69) and a trend in the SESSION factor (F_(1,23)_ = 4.14, p = .054, η^2^ = 0.152, BF = 1.56). Together, these effects revealed an increasing beta-band power in frontal areas that peaked around the middle of the task (BIN 1 vs BIN 5: p = .002, BF = 35.56; BIN 1 vs BIN 10: p = .404, BF = 1.63, Bonferroni correction for ten comparisons was applied to the p-value) (Fig 2B), with the highest amplitude in the S12 session (Fig 2B). We found no effect in the BIN-SESSION interaction (p = .12, BF = .25).

### Beta-band specificity

Since some of our results appear at odds with the current physical fatigue literature, we checked if they were persistent when considering more classical, absolute beta-bands, specifically Low-Beta (Lβ, 13-21 Hz) and High-Beta (Hβ, 21-31 Hz). We first ran RM-ANOVAs with BIN (1 to 10) and SESSION (S10, S12) as fixed factors on Lβ and Hβ Cz quantified during contractions. While only a BIN-SESSION interaction effect was found when considering Lβ Cz (F_(9,207)_ = 2.86, p = .003, η^2^ = 0.11, BF = 7.69) (p > .12, BF < .80, for BIN and SESSION), Hβ Cz revealed a very strong BIN effect (F_(9,207)_ = 4.92, p < .001, η^2^ = 0.18, BF = 3493.17) (p > .37, BF < .37, for SESSION and BIN-SESSION interaction), indicating a decreasing power over the task (Fig 3A). Whereas Lβ does not replicate Iβ dynamic, Hβ aligns thoroughly with this latter, thus supporting our findings and emphasizing a Hβ-band specificity to the power modulations associated with lower limbs contractions during physical fatigue (Neuper & Pfurtscheller, 2001b; Pfurtscheller et al., 2000).

**Figure 3.**
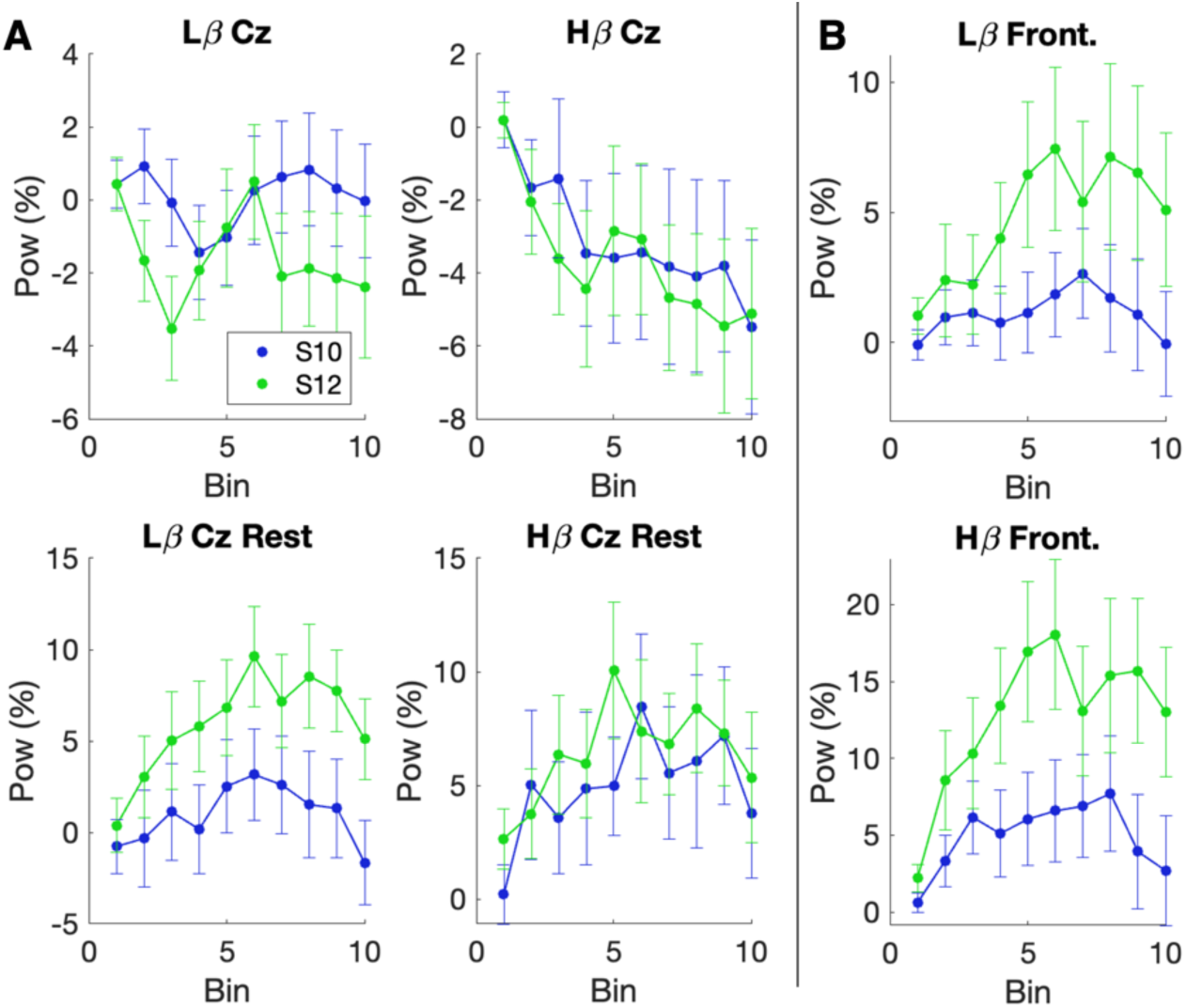
Beta-band power modulations in low and high-beta bands. A. Motor areas. All panels represent the relative power changes over the 100 contractions (respective to the first contractions, see Methods for details) quantified in Cz in Low-Beta (Lβ) and High-Beta (Hβ) during contractions (panels 1 and 2) and at rest (panels 3 and 4). B. Frontal areas. The panels display Lβ and Hβ power relative changes over the 100 contractions, quantified in frontal electrodes during contractions. For A and B, the dots represent the averaged distribution, and the error bars the SEM.

We then ran the same analysis on Lβ and Hβ Cz quantified at rest. We found a BIN effect for both Lβ (F_(9,207)_ = 3.88, p < .001, η^2^ = 0.15, BF = 60.66) and Hβ (F_(9,207)_ = 3.26, p < .001, η^2^ = 0.12, BF = 5.92), revealing an increasing power over the task (Fig 3A). No SESSION or BIN-SESSION interaction effect was found for both variables (p > .088, BF < 1.17). These results align with our findings and suggest that the increasing power observed, which promotes maintaining the resting state, is not restricted to sub-beta bands.

Finally, we performed the same RM-ANOVAs considering frontal electrodes in Lβ and Hβ during contractions. We found a very mild BIN effect (F_(9,207)_ = 2.13, p = .028, η^2^ = 0.085, BF = 0.50) for Lβ, revealing an increasing power over the task. There was no SESSION or BIN-SESSION interaction effect (p > .076, BF < 1.28). Regarding Hβ, we found a very strong BIN (F_(9,207)_ = 4.49, p < .001, η^2^ = 0.16, BF = 428.31) and a SESSION effect (F_(1,207)_ = 4.74, p = .040, η^2^ = 0.17, BF = 2.01), showing an increasing power over the task, with the highest amplitude in the S12 session (Fig 3B). There was no BIN-SESSION interaction effect (p > .087, BF < 0.33).

Overall, these results corroborate the effects on Iβ and highlight the importance of considering narrower beta-bands when investigating physical fatigue (Kilavik et al., 2013). While the pattern observed at rest in Cz was broad, the modulations associated with contractions were specific to Iβ and Hβ.

### Connectivity over physical fatigue

After investigating the changes in beta-band power in motor and frontal areas, we explored the connectivity between these regions as well as with the EMG during the task. Specifically, we first investigated the connectivity via subject-by-subject correlations between Iβ in motor areas (i.e., Cz electrode) and EMG and between Iβ in motor and frontal electrodes. Our results reveal that the correlation coefficients across participants were negative between Iβ Cz and EMG indexes in both sessions (S10: t_(23)_ = 2.34, p = .028, d = 0.48, BF = 2.05; S12: t_(23)_ = 2.47, p = .021, d = 0.50, BF = 2.57), while we found a positive correlation between Iβ Cz and Iβ in frontal electrodes (S10: t_(23)_ = 7.23, p < .001, d = 1.48, BF = 6.80×10^+4^; S12: t_(23)_ = 2.93, p = .007, d = 0.60, BF = 6.20) (Fig 4A). Overall, the antagonistic relation between EMG and Iβ Cz supports the assumption that both measures reflect increased MC activation: the enhanced EMG activity reflects the increasing central motor command compensating for the growing muscle fatigue, which arises from the heightened MC activation, underlined by the decreasing beta-band power in Cz.

To deepen our understanding of how physical fatigue modulates the interaction between the motor and frontal areas, we conducted Granger Causality (GC) analyses during the contractions between these regions. We performed RM-ANOVA with BIN (1,2,3,4,5, each BIN comprising 20 contractions) and SESSION (S10, S12) as fixed factors, on the GC values associated with the two directions flow, namely Cz → Front. and Front. → Cz, both in the temporal and spectral domain. A substantial BIN effect was shown from frontal electrodes to Cz in the temporal domain (F_(4,92)_ = 4.29, p = .003, η^2^ = 0.16, BF = 7.95), while an anecdotal one was shown in the opposite direction (i.e., Cz → Front.) (F_(4,92)_ = 2.52, p = .047, η^2^ = 0.099, BF = 0.74) (Fig 4B), both supporting an increased communication between motor and frontal areas as fatigue rises (Jiang et al., 2012). Regarding the spectral domain, we investigated specifically the Iβ frequency, as we found significant modulations of Iβ oscillations both in the motor and frontal regions during physical fatigue. However, the GC analysis revealed no effects (p > .93, BF < 0.52, both directions).

**Figure 4.**
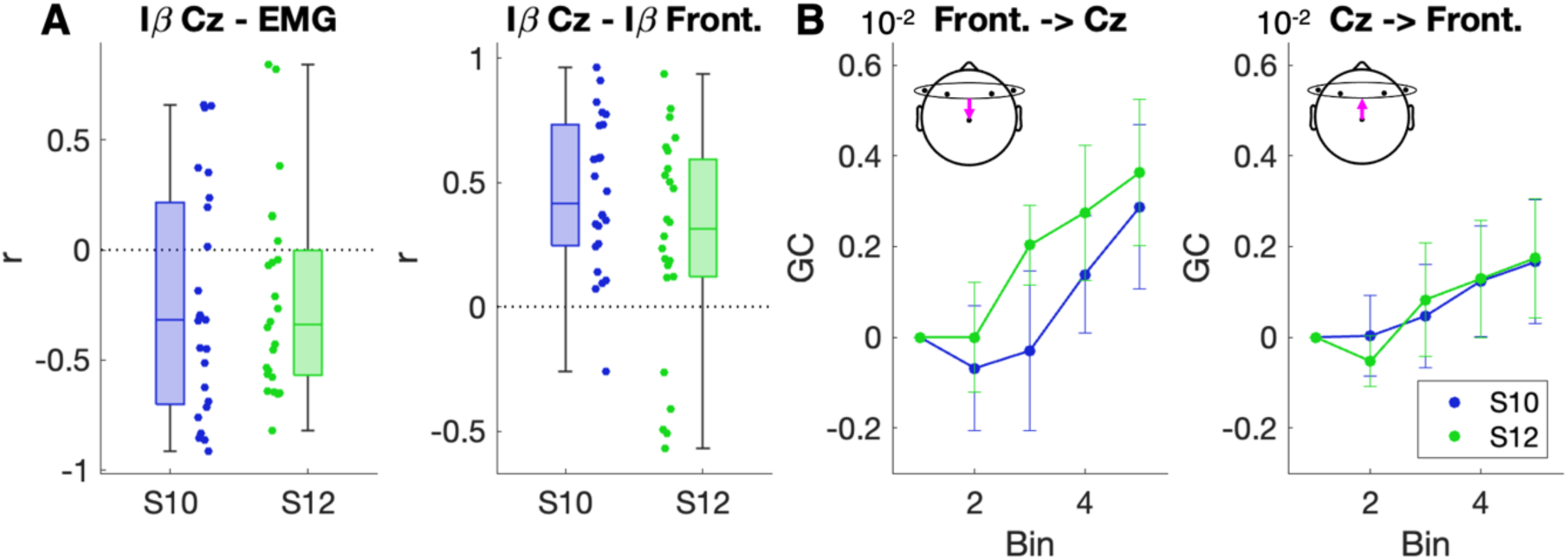
Functional and effective connectivity. A. Correlations. Within subjects’ correlations were performed between Iβ Cz and EMG, and Iβ Cz and IB Frontal, for each session independently. The resulting r values were then compared using t-tests. B. Granger causality (GC). GC was performed to assess the relative changes in the bidirectional information flow between frontal and motor areas. Each bin results from the integration of 20 contractions. The dots represent the averaged distribution, and the error bars the SEM. The first bin is 0 centered for both sessions.

### Fatigue-related power modulations during resting states

After investigating the ongoing changes during the task, we investigated power variation rates between pre- and post-task resting states for Iβ in motor and frontal regions. This analysis aimed to explore how fatigue affects the resting state beta-band power, which might serve as an index of the strength of the status quo maintenance. While no effect was found in the frontal areas (p > .46, BF < 0.28, for both sessions), we observed a significant increase in Iβ power in Cz for both sessions (S10: p = .001, d = 0.77, BF = 34.36; S12: p < .001, d = 0.90, BF = 148.21) (Fig 2D). However, pairwise comparison performed on Iβ power in Cz between both sessions did not reveal a significant difference (p = .103, BF = .81, unilateral pairwise t-tests, H1: S10 < S12). Note that the Iβ power increase observed in Cz was also found when considering Lβ (p < .016, d > 0.53, BF > 3.34, for both sessions) and Hβ (p < .001, d > 0.86, BF > 92.89, for both sessions).

Since enhanced beta-band power promotes the preservation of the current motor set (Engel & Fries, 2010), we hypothesized that the greater the beta-band power during resting states, the larger the desynchronization due to the contraction (Fig 5A). To test this hypothesis, we split the participants into two groups based on the median Iβ power resting state pre-post variation rate and tested whether the Iβ Cz decrease during the task was different between them. Consistently with our hypothesis, we found a significant effect in S10 and a trend in S12 (S10: t_(22)_ = –2.35, p = .014, d = 0.96, BF = 4.74: S12: t_(22)_ = –1.60, p = .065, d = 0.64, BF = 1.61, unilateral independent samples t-test, H1: Low Iβ Post < High Iβ Post), highlighting a larger desynchronization over the task for the ‘High Iβ Post’ group, compared to the ‘Low Iβ Post’ one (Fig 5B). To further corroborate this effect, we tested the same hypothesis with a different approach. We computed the correlation between the pre-to post-variation rate in Iβ power in Cz during the resting state and the slope of Iβ Cz during the task, a variable that allows us to assess the trend of Iβ power over the task. As hypothesized, the increase in Iβ power over Cz during rest was negatively correlated with the Iβ Cz decrease during the contractions in both sessions (S10: r = –0.53, p = .004, BF = 13.81; S12: r = –0.37, p = .038, BF = 2.12, unilateral test, H1: negative correlation): in other words, the larger the increase in beta power at rest after the task, the larger the desynchronization during the contractions (Fig 5C).

**Figure 5.**
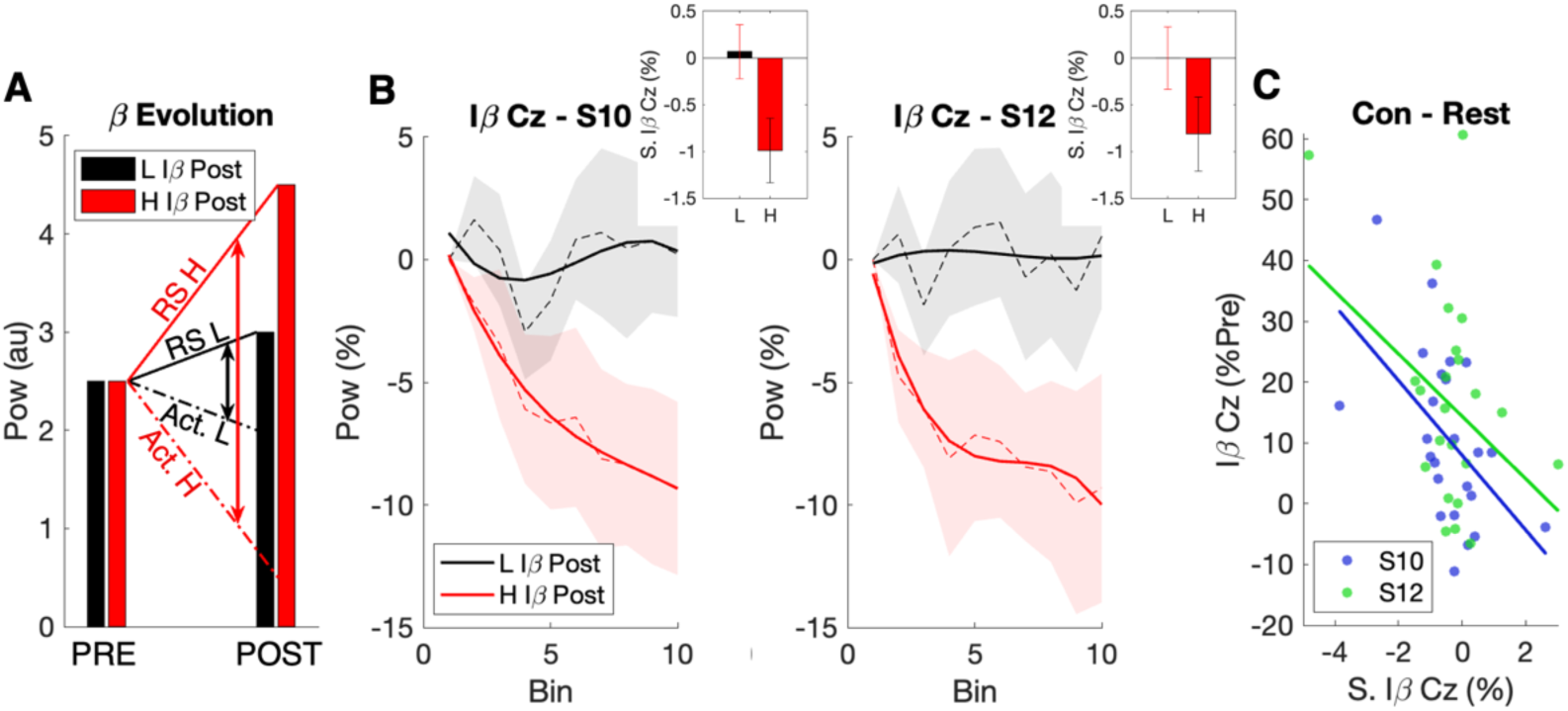
Relation between beta-band modulations during resting-state and in contraction. A. Schematic representation of Beta evolution. ‘RS’ stands for Resting-State (Baseline power). ‘Act.’ stands for activation, which reflects the necessary activation threshold (beta desynchronization threshold) needed to perform the contraction at the required intensity. The double vertical arrows represent the amount of desynchronization for each group. Considering two hypothetical groups that have different (low (L) or high (H)) relative changes between the Pre and Post RS beta-band power, we might expect to find a larger desynchronization for the H group. In other words, the larger the increase in beta-band power RS after the task, the larger the desynchronization during the contractions. B. Iβ Cz over bins for each group and each session. The participants were divided into two groups (L Iβ Post and H Iβ Post) according to a median split performed on the Iβ power RS pre-post variation rate. Then, we represented the Iβ power evolution during contractions for these two groups. The dashed line represents the participants’ average, the shaded area of the SEM, and the solid line, a polynomial fit with 3 degrees of the average. Each group’s first bin is zero-centered. The bar plots represent the slope of the Iβ Cz power computed over bins for each group. C. Correlations between Iβ Cz slope (S.) in contraction and Iβ Cz power pre-post task variation rate.

### Modelling beta-band power fatigue modulations

Somewhat contrary to previous studies that suggested an increase of beta power during contractions due to fatigue, our experimental data revealed that beta-band oscillations in the MC decrease over submaximal contractions due to physical fatigue (Fig 2B), supposedly driven by some top-down modulatory effect (Fig 4B). To understand such an effect, we implemented a simple but biologically plausible model composed of 1000 Izhikevic neurons (Izhikevich, 2003). Specifically, the network was composed of excitatory and inhibitory neurons (Fig 6A), whose baseline activity spontaneously generated bursts of beta-band oscillations. To simulate the effect of a motor contraction, 20% of the excitatory neurons received a top-down inhibitory modulation at 1500 ms (Fig 6B) from frontal regions (as suggested by our experimental results – Fig 4B). Despite its simplicity, the network successfully replicates the beta-band desynchronization (Fig 6A, 6B). Next, in order to replicate the fatigue effect, we linearly increase such inhibitory modulation over five repetitions, and we run the simulations 10 times. To statistically test such fatigue effect driven by the top-down inhibition, we performed an RM-ANOVA on the beta-band power changes with BIN (1 to 5) as a fixed factor. We found a strong BIN effect (F_(4,36)_ = 33.71, p = < .001, η^2^ = 0.79, BF = 8.59×10^+8^), revealing an increase in the beta-band desynchronization over the bins due to a rise in the inhibitory input (Fig 6A, 6C). These results, in line with our experimental data (Fig 2B), provide a simple yet plausible implementation of such beta-band desynchronization during the contraction, yielding a mechanistic explanation of our experimental observations.

**Figure 6.**
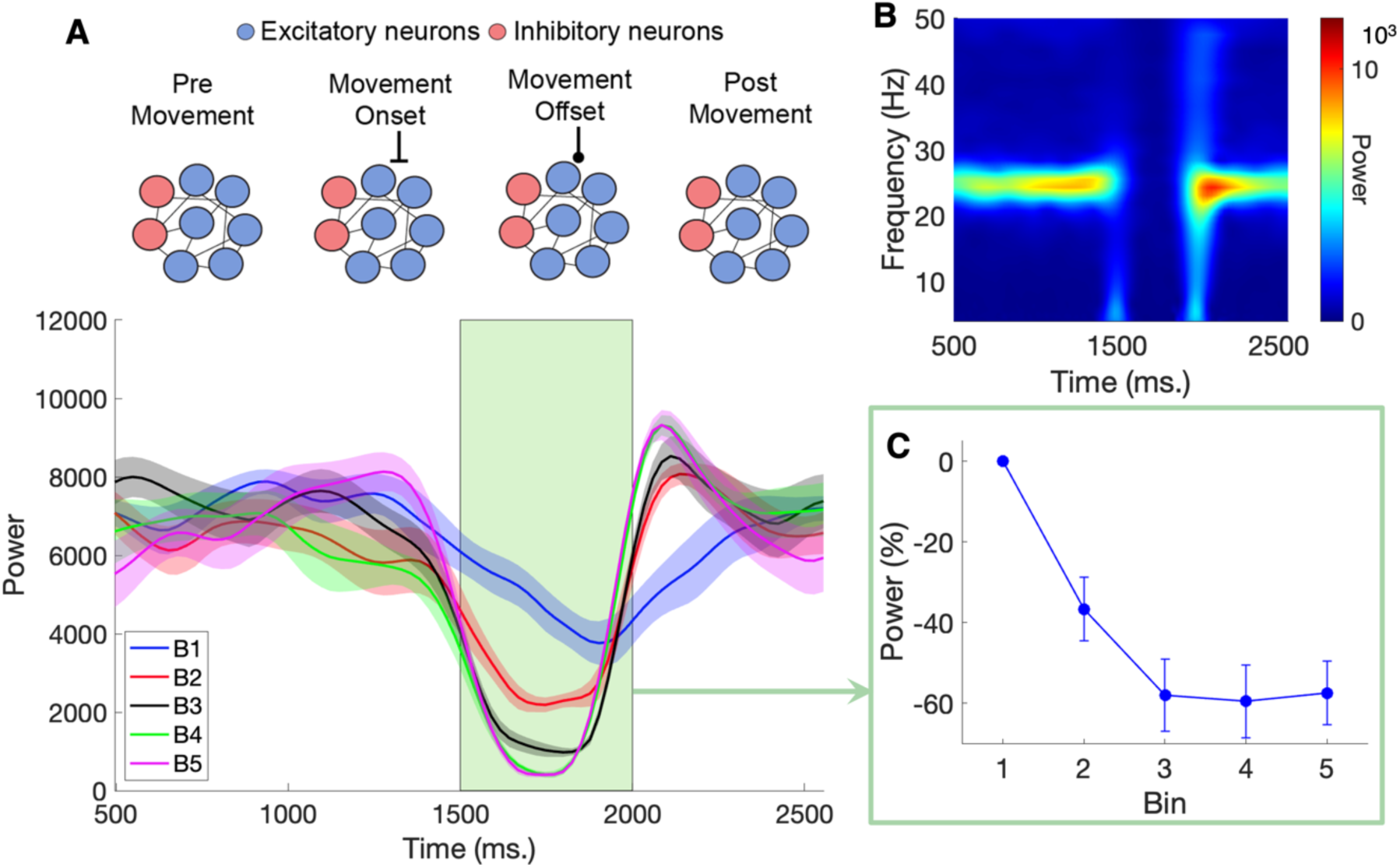
Spiking network model of beta-band desynchronization during physical fatigue. A. Spiking network and beta-band modulations. The upper part of the panel illustrates the 1000 interconnected Izhikevich neurons, 80% excitatory and 20% inhibitory. The ‘movement’ period corresponds to an inhibitory modulation sent for 500 ms to a subset of excitatory neurons. The lower part of the panel displays the beta-band power generated by the spiking network and modulated over time. A desynchronization occurs during the inhibitory input (green area). The five Bins (B) represent an incremental inhibitory input (from Bin 1 to Bin 5) that simulates an increasing signal from the frontal to the motor areas as fatigue rises. B. Typical time-frequency map. The network generates beta power centered around 25 Hz, followed by a desynchronization and a slight rebound at the end of the movement. C. Beta-band power modulations. This panel represents the relative power changes quantified during the ‘movement’ period over Bins. The dots represent the averaged distribution of the 10 simulations, and the error bars the SEM.

## III. Discussion

In this study, we proposed a novel experimental design whose findings reconcile the role of beta-band oscillations during physical fatigue within the status quo framework (Engel & Fries, 2010). In particular, the EEG recordings over the motor areas revealed a decrease in beta-band power during the contractions but an overall increase in the resting periods, during and after the task. We also reported a rise in beta-band power in the frontal regions during contractions. Interestingly, motor and frontal oscillatory dynamics modulations during contractions were underlined by an increased information flow during physical fatigue, mainly from the frontal to the motor areas. Building on these results, we propose a unifying explanation that describes the functional role of beta-band oscillations during physical fatigue within the status quo theory.

### Beta-band power modulations in motor areas during physical fatigue

Our findings highlight two distinct beta-band power dynamics in motor areas during physical fatigue. On the one hand, we found a reduction of beta-band power during contractions, deviating from most of the current physical fatigue literature, which reports the opposite effect (Bailey et al., 2008; Ciria et al., 2018; Enders et al., 2016; Hosang et al., 2022; Kubitz & Mott, 1996; Lin et al., 2021; Suviseshamuthu et al., 2021; Yang et al., 2009). On the other hand, our results revealed a beta-band power increase at rest, in line with previous findings (Crabbe & Dishman, 2004). Remarkably, both dynamics aligned thoroughly with the status quo interpretation (Engel & Fries, 2010). The decreasing beta-band power during contractions reflects a change (i.e., increase) in MC activation necessary to sustain a submaximal task at a constant intensity through physical fatigue (Taylor et al., 2016). On the other hand, the heightened beta-band power at rest supports the maintenance of the resting state. Although our study places the physical fatigue paradigm within the status quo theory, our results differ from the physical fatigue literature, which reports a rise in beta-band power during contractions with fatigue, suggesting that it reflects an increase in MC activation (Enders et al., 2016; Kubitz & Mott, 1996; Lin et al., 2021; Yang et al., 2009). In the following, we propose an alternative explanation that may reconcile previous findings with the status quo theory and our results.

### Reconciling the fatigue literature with the status quo theory

The rise in the MC activation that occurs during a submaximal exercise reaches a plateau before exhaustion, as shown previously using fMRI (Liu et al., 2003) and as suggested in EEG studies investigating the beta-desynchronization amplitude in relation to the contraction intensity (Dal Maso et al., 2012; Stancák et al., 1997). When this activation ceiling is reached, the task can still be performed, but the central fatigue – the decreasing ability of the central nervous system to recruit the motor units (Bigland-Ritchie, 1981) – continues to grow. To counter this effect in advanced stages of fatigue, alternative brain areas supplement the MC function to contribute to the muscle output (Liu et al., 2007; Yang et al., 2009), enabling task continuation while limiting the development of central fatigue. Therefore, one could speculate that the temporal dynamics of beta-band power over a fatiguing task follows a three stages pattern: (i) a first decrease to initially cope with both muscle and central fatigue, (ii) a stabilization of the activity as the entire pool of available neurons is recruited, and (iii) a final increase with the rising central fatigue, with some of the motor command reallocated to other brain areas. Considering our experimental design, our task may have tapped into the first two stages, whereas other tasks performed in the literature – especially those conducted to exhaustion, may have reached the latest stage. Further studies will be necessary to shed light on the temporal dynamic of beta-band power during a fatigue task to exhaustion.

### A push-pull effect? Relation between beta-band power at rest and in contractions

Our results revealed a significant increase in beta-band power recorded at rest in motor areas, both during the task (between contractions) and after the task (3 minutes resting-state period). Since an increase in beta-band power after a contraction is assumed to reflect the inhibition of the sensorimotor cortex (Neuper & Pfurtscheller, 2001a; Pfurtscheller et al., 1996; Sallard et al., 2014; Solis-Escalante et al., 2012), the growing beta power at rest may represent an inhibitory mechanism of the motor system in response to the beta power decrease observed during contractions. Specifically, we propose that an increase in beta-band power may reflect the rising central fatigue at the supra-spinal level (Gandevia, 2001), thus making the release of inhibition to initiate a new contraction (and, per se, the effort continuation) harder (Barone & Rossiter, 2021). This interpretation aligns with the high resting-state beta-band power observed in aging (Rossiter et al., 2014) and pathological conditions such as Parkinson’s disease (Jenkinson & Brown, 2011; Little & Brown, 2014; Oswal et al., 2013) that is directly related to movement execution impairment. In addition, transcranial alternative current stimulation studies have indicated decreased motor performance (Joundi et al., 2012; Pogosyan et al., 2009; Wach et al., 2013) and increased fatigue (both physical and subjective) (Rönnefarth et al., 2017) when enhancing beta synchronization, supporting the hypothesis that heightened beta oscillations make the task continuation harder. Additionally, our data showed that participants with the greatest increase in beta-band power at rest exhibit the largest beta desynchronization during contractions, corroborating the presence of such a compensatory mechanism. All in all,the functional role of beta-band oscillations in physical fatigue seems to encompass two processes. On the one hand, there is an increasing synchronization at rest with physical fatigue, which might indicate the supra-spinal level of central fatigue that inhibits the system for subsequent movements. On the other hand, this inhibition is countered by a larger desynchronization required to sustain effort at the necessary intensity, hence producing a ‘push-pull’ effect.

### Function role of beta oscillations in the prefrontal cortex

Beta power quantified during contractions over frontal regions revealed an increase during the task, peaking around the midpoint of the task. While this enhanced beta-power is in line with previous studies (Crabbe & Dishman, 2004; Hosang et al., 2022), its functional role in physical fatigue remains poorly understood. During physical effort, the PFC plays a crucial role in integrating physiological signals and determining the subsequent decision to continue (or not) the effort (McMorris et al., 2018; Robertson & Marino, 2016). As beta oscillations in the PFC have been proposed to reflect top-down cognitive control (Buschman & Miller, 2007; Stoll et al., 2016) and especially to act as a filter against distractors (Schmidt et al., 2019), it may serve a similar function during physical effort. One hypothesis is that enhanced beta-band power could actively protect the internal goal (i.e., continuing the task) over rising uncomfortable sensations arising from physical fatigue. Interestingly, a recent study suggested that the subjectively perceived hardest period of a fatiguing task with a known duration occurs between ∼50 and 75% of its duration rather than increasing linearly over the task (Camparo et al., 2022). These findings seem to align with the dynamics of the beta-band power in frontal areas in our data, which reveal an inverted U-shape peaking around the middle of the task. This interpretation supports the compelling hypothesis that beta-band power in frontal areas reflects a neural correlate of the cognitive control needed to pursue a physical effort through fatigue.

### Increased bidirectional communication through physical fatigue

We found an increasing bidirectional communication between motor and frontal areas through physical fatigue, mainly from frontal to motor areas. To the best of our knowledge, no EEG study has looked at this specific relationship before. Regarding prominent models of physical fatigue (McMorris et al., 2018; Tanaka & Watanabe, 2012), the increasing flow from frontal to motor areas could reflect a growing signal from the PFC to the MC to continue the task despite the occurring fatigue. One previous fMRI study has shown similar results when comparing connectivity through cross-correlation between the two areas in a high versus low fatigue stage (Jiang et al., 2012). Hence, the increasing input from the PFC could be the required signal that allows releasing the increasing beta-band power active inhibition at rest, leading to larger desynchronization through physical fatigue, as shown in our experimental data and model. Conversely, the minor increase in the flow from MC to PFC areas may be interpreted as a form of corollary discharge (Bigliassi, 2015; Poulet & Hedwig, 2006) that allows adjusting the PFC output regarding the actual motor command through an indirect cortico-basal-ganglia-thalamo-cortical loop (Tanaka & Watanabe, 2012).

### Conclusion

Our study shed light on the functional role of beta oscillations during physical fatigue by integrating it within the status quo theory. We identified two concurrent mechanisms in the motor areas as fatigue rises: a decrease in beta-power during contractions and an increase in rest periods (during and after the task). While the latter promotes the maintenance of the resting state, the former reflects the increase in the MC activation necessary to cope with muscle fatigue, driven by an increasing signal from the frontal areas. By integrating the physical fatigue paradigm within the status quo theory, we further proposed a unifying explanation that may reconcile conflicting evidence in the literature. Finally, it is important to consider that in this study, we investigated the functional role of beta oscillations in physical fatigue through an analytic quadriceps isometric task. Further studies will generalize our findings to a more ecological context (e.g., whole-body exercise), accounting for other relevant physiological parameters (muscle typology, cardiovascular capacity, etc.).

## IV. Methods

### 1. Participants

Twenty-eight participants took part in this study (16 women, age: 24 ± 4, weight: 70 ± 18, mean ± SD). All participants had no known history of neurological diseases or physical injuries and practiced physical activity once to several times a week. Of the 28 participants, 24 could perform the whole protocol and were retained for the analyses. The study was approved by the French Ethical ‘Comité de Protection des Personnes’ (number: 2023-A00412-43). All individuals provided their informed consent.

### 2. Data Acquisition

All participants were seated during each session on an isokinetic ergometer (Biodex) with both knees and hips flexed at 90°. Their right leg was firmly fixed to the ergometer accessory at the ankle, just above the malleolus, with a non-compliant strap. A minimization of their body movements and the contribution of other muscles (aside from the quadriceps) were ensured through two shoulder harnesses and an abdominal one. Installation settings were optimized for every subject individually and kept identical within subjects for both days.

Torque and electromyography (EMG) were recorded at 2000 Hz using a Biopac MP150 system and AcqKnowledge software. EMG was recorded through three pairs of electrodes (AG-Cl, diameter: 11 mm, interelectrode distance: 2 cm), which were placed on the Vastus Medialis, Vastus Lateralis, and Rectus Femoris according to SENIAM recommendations. Before placement, the skin was carefully prepared by shaving and applying alcohol to ensure low impedance (<5 KΩ). The reference electrode was positioned on the patella of the contralateral knee. EEG was recorded using a 64 electrodes Biosemi Active 2 system with 64 Ag-AgCl sintered active electrodes at 2048 Hz. The EEG electrode locations followed the 10–20 international system. The offset was kept below 20 mV using an active gel. All acquisitions were performed with the light off to minimize electrical noise.

The main task was programmed in Python and displayed through a Windows computer (Resolution: 1920 × 1080; Refresh rate: 60 Hz), using a NiDAQ acquisition card (NI USB-6218, Sample rate: 2000 Hz) to read the torque data (low-pass filtered at 10 Hz) in real-time. The same computer also generated the triggers used to mark the onset and offset of each contraction towards Biopac and Biosemi, allowing us to synchronize all our signals offline.

### 3. Experimental Protocol

#### 3.1. Procedure

All participants came to the lab on four days (spaced 3 to 10 days apart) to perform four sessions. In the current paper, we focus only on two of these days/sessions and do not mention the two other ones further, which are part of a larger protocol. On each day, the participants performed the following steps (Fig 1A):

- Warm-up (consisting of 15-20 contractions with increasing intensity, approximately from 20 to 80% of their estimated Maximal Voluntary Torque (MVT))
- 2 Maximal Voluntary Contractions (MVC) (3-4 s each, 30 s spaced)
- 1 MVC, 20 s of rest, 1 Long MVC (30 s), 30 s of rest, 3 minutes of EEG recording at rest (eyes opened)
- 5 minutes of rest
- Main task: 100 contractions (see below)
- 1 MVC, 20 s of rest, 1 Long MVC (30 s), 30 s of rest, 3 minutes of EEG recording at rest (eyes opened)

Depending on the session, the main task consisted of 100 contractions of 10 or 12 seconds each. For both sessions, each contraction was preceded by a preparation time of 0.75 s (when the participants tried to reach the targeted force) and was separated by a 5 s resting time. In practice, a sound indicated to the participants when to begin and end each contraction. Concomitantly with the starting sound, a rectangle was displayed in the middle of a black screen to provide online feedback to the participants on their actual force compared to the individualized (see below) targeted one. This procedure was achieved by updating the rectangle’s color online according to the ongoing force produced by the participants: red indicated that participants were below the targeted force, blue above, and green in the aimed range. The green range spanned from -5% to +5% around the targeted force. 0.75 s after the start of the contraction, a digital clock was displayed in the middle of the rectangle to inform participants of the time of their contraction. After the ending sound, the clock (10.0 or 12.0 s) remained on the screen for 1 s, then the number of contractions already performed was displayed during the resting time.

#### 3.2. Individualized torque target determination

We determined the targeted torque for each participant to generate enough physical fatigue during the task while allowing participants to finish it without excessive trembling (to minimize muscle artifacts). Accordingly, we chose an absolute fixed load of 40 N.m following pilot experiments, corresponding to ∼10 to 20% of the MVC of most participants, allowing them to perform all of the 100 contractions. As 40 N.m constitutes a potential ‘heavy’ load (>20%) for participants with an MVT under 200 N.m, we decided to set their targeted force at 20% of their MVC. Notably, the chosen load was kept identical for the two sessions for the same individual, allowing sessions to be compared. Despite these criteria, four of the participants included were not able to perform the 100 contractions at the required intensity and were removed from any further analyses.

### 4. Data pre-processing

All data pre-processing and analyses were performed offline using custom MATLAB (version 2022b) scripts and EEGLAB functions (Delorme & Makeig, 2004).

Torque and EMG data were down-sampled offline at 1000 Hz. Torque data were low-pass filtered at 10 Hz, EMG data were band-pass filtered between 20 and 500 Hz, and notch-pass filtered between 47 and 53 Hz.

EEG data were down sample offline at 250Hz. A notch filter, from 47 to 53 Hz, and a high-pass filter, at a cut-off frequency of 1 Hz, were applied. Channels with excessive noise were automatically rejected using the CleanRawData plugin set up with a 5 s ‘flat line criterion’, a 0.6 ‘minimum channel correlation’, and a kurtosis superior to five times the mean. On average, 61.8 ± 1.1 (mean ± SD) channels were kept for the analysis. Then, electrodes were re-referenced following a common average reference. An Independent Component Analysis was performed using the Infomax algorithm. The resulting components were labeled using the ICLabel plugin (Pion-Tonachini et al., 2019). An automated rejection of all components labeled as muscle, eye, heart, line noise, channel noise, or other, with a probability above 90%, was applied. On average, 12.5 ± 4.5 components were rejected. The previously removed channels were then interpolated.

The EEG data recorded during the two resting state periods were epoched into 2 s non-overlapping windows. The EEG and EMG data from the main task were segmented in epochs of 18 s, each starting 3 seconds before the onset of the contraction (as determined by its trigger) and ending 15 seconds after it. A last epoch’s rejection procedure was applied to the EEG data recorded during the main task to remove additional noise. For each subject and electrode, epochs were rejected with a standard deviation superior to 1.5 times the interquartile range (IQR) over the 3rd quantile or inferior to 1.5 times the IQR below the 1st quantile. This procedure removed 4.3 % and 3.5 % of the epochs in the S10 and S12 sessions, respectively.

### 5. Data analysis

#### 5.1. Force and EMG

<u>Pre-Post.</u> The maximal torque among the three MVCs before the task was retained to quantify the pre-test Maximal Voluntary Torque (MVT) measurement. The post-test measurement corresponded to the maximal torque recorded during the post-MVC. The ability to sustain an MVC was quantified as the relative difference (labeled ‘Long Max Diff.’) between the maximal torque reached during the five first seconds and the five last seconds of the long MVC (Lebesque et al., 2022). Torque integral (‘Long Max Int.’) was also extracted to quantify thoroughly the force-sustaining ability. The variation rate ((post – pre)/pre *100) was computed for each variable.

<u>Main-Task.</u> EMG was quantified as the root-mean-square (RMS) during each contraction of the main task throughout 2 to 10 s (0 s being the onset of the contraction). Importantly, this period allows us to (i) avoid taking into account the beginning of the contraction, where the participants’ torque is not stabilized yet, (ii) avoid taking into account the end of the contraction in the S10 session (as participants may anticipate it), (iii) have an identical time of comparison for the two sessions (S10 and S12). In this period of interest, the EMG was specifically quantified as the RMS over 180ms (non-overlapping) windows, which were normalized by the RMS EMG associated with the previously selected maximal torque (centered from -90 ms to +90 ms around it) of the respective muscle and session. The three muscles were then averaged over time, yielding one EMG value associated with each contraction in the main task. Lastly, these EMG values were averaged in bins of 10.

#### 5.2. EEG Spectral Power

Spectral power was obtained by applying a continuous wavelet transform on the pre-processed epoched EEG data for the frequency range from 1 to 50 Hz. The length of the wavelets was linearly increased from one cycle at 1 Hz to twenty cycles at 50 Hz to optimize the tradeoff between temporal and frequency resolution on the whole frequency range. Removed contractions were then linearly interpolated.

Spectral power of Low-Beta (Lβ, 13-21 Hz) and High-Beta (Hβ, 21-31 Hz) bands was extracted. We also defined an Individualized Beta (Iβ) band for each subject (Fig 1B) to account for the inter-individual variability in the beta desynchronization-specific frequency band. To this end, we normalized the raw spectral power during the period of interest (from 2 to 10 s, see above) to the rest (-3 to -1 s) in the Cz electrode for each contraction (Pfurtscheller & Lopes da Silva, 1999). The median of all contractions (100) was computed for each session, providing two time-frequency maps for each subject, which were then averaged together. From this individual time-frequency map, the characteristic movement-related beta desynchronization was quantified on a wide beta band (13-40 Hz), whose bounds were set to cover the broad range of desynchronization frequencies observed across participants. Over these frequencies and time range, the individualized frequency was retained as the one with the largest desynchronization for each subject respectively. Iβ was finally defined as ranging from – 2 to + 2 Hz around this frequency. The distribution of the Iβ frequencies was comprised between 17 and 36 Hz (Fig 1C), consistent with previous work showing a higher frequency specificity associated with lower limb contraction (Neuper & Pfurtscheller, 2001b; Pfurtscheller et al., 2000).

Power from the lateral-frontal (F7-F8) and mid-frontal (F3-F4) electrodes were averaged to constitute a ‘Frontal’ cluster. These electrodes were chosen as they are thought to reflect the activity of the ventrolateral and dorsolateral prefrontal cortex (Okamoto et al., 2004; Robertson & Marino, 2015). Cz was retained as reflecting the MC activity associated with the quadriceps contraction (Dal Maso et al., 2012; Forrester et al., 2006), a location confirmed by visualizing topographically the beta desynchronization.

<u>Pre-Post.</u> Raw spectral powers from pre- and post-EEG resting state recordings were independently averaged over epochs (90) and time (0 to 2 s). The variation rate was computed between the pre and post-value for each frequency band and region of interest.

<u>Main Task.</u> For each contraction, raw spectral power bands on the period of interest (2 to 10 s) were normalized by the corresponding session’s first contractions – on the same 2 to 10 s period averaged over time, allowing to focus on the fatigue effect over the contractions. Specifically, the normalization consisted of subtracting and dividing by the baseline and multiplying by 100, thus leading to the expression of power values as a percentage of a non-fatigued state. The baseline consisted of contractions 2 to 5, which were averaged together. Note that the first contraction was not considered for this normalization as its torque presented substantial variability for both sessions. The resulting values were averaged over time and frequencies, leading to one power value by frequency band, contraction, and electrodes. Following the same procedure, we also computed Lβ, Hβ, and Iβ in Cz at rest, that is, on a period of interest from -3 to -1 s regarding the start of the contraction. Finally, we averaged over contractions in bins of 10 contractions to further increase the signal-to-noise ratio.

<u>Within contraction.</u> For each contraction, Iβ power in Cz in the period of interest (from 2 to 10 s) was normalized to the rest period (-3 to -1 s). The average of the Iβ power in Cz was then performed over the 100 contractions to obtain one typical contraction.

#### 5.3. EEG Connectivity

To investigate the relationship between the motor and frontal regions, as well as between the motor areas and the EMG, we performed Pearson’s correlation between Iβ Cz and the EMG, and between Iβ Cz and Iβ Frontal, for each subject respectively and then used a bilateral one-sample t-test to test whether the r values significantly differed from zero.

The evolution of the directed information flow between motor and frontal regions over the task was assessed using Granger Causality (GC), both in the temporal and spectral domain (at Iβ frequency), computed through the MVGC toolbox (Barnett & Seth, 2014). We considered the 2 to 10 s period of interest within each contraction and divided it into eight 1 s epochs to approximate the stationary signal requirement (Seth et al., 2015). Contractions were then binned by 20, leading to integrating ∼160 epochs (20*8, minus the potentially removed contractions (3.9% of all the trials)) to reconstruct one GC value. A model order of 17 was chosen according to the Akaike Information Criterion and Bayesian Information Criterion averaged over all participants, and the maximum autocovariance lags were fixed at 1500. This analysis was run between the pair of electrodes: ‘Cz – F3’, ‘Cz – F4’, ‘Cz – F7’, and ‘Cz – F8’, from which the resulting GC values were averaged together, providing a measure of directionality between motor areas (‘Cz’) and frontal areas. For both the temporal and spectral domains, the first bin of both sessions was centered in 0 to account only for the relative changes.

### 6. Modelling

We performed computational simulations to model the fatigue effect on beta-band oscillations. We implemented a neuronal network composed of 800 excitatory and 200 inhibitory neurons to represent the beta-band activity in the MC. Each neuron was modeled by the biologically plausible dynamics proposed by Izhikevich et al. (2004). Specifically, the membrane potential *v* was defined as:

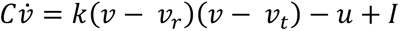

Where u represented the recovery current:

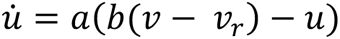

When the membrane potential reached the threshold *v_t_*, both the current and the membrane potential were set to: *v* = *c* and *u* = *u* + *d*. Importantly, *a, b, c, d*, and *k* were model parameters determining the neurons’ dynamics and firing rate. We set the neurons’ parameters to generate beta-band oscillations at rest (i.e., without external stimulation), having similar values as in the Resting State (RS) proposed in Izhikevich (2003). Specifically, we set *a* = 0.04, *b* = 0.3, *c* = −65, *d* = .25, *k* = 0.015, *C* = 4, *v_t_* = −25 and *v_t_* = −85. After one second at rest, we model the onset of a motor contraction by providing 20% of the excitatory neurons with a top-down inhibitory modulation from frontal regions. We model the effect of fatigue by producing five contractions with a linearly increase in the top-down modulation, and we run the simulations 10 times. We assessed the effect on the network dynamic by computing the beta-band power over time, that is, before, during, and after the motor contraction. We eventually compared the beta-band desynchronization as a function of the top-down modulation strength.

### 7. Statistics

All the statistical tests were performed using both the frequentist and the Bayesian frameworks to account for the acceptance of the null hypothesis (Masson, 2011). Significance in the frequentist approach was set at p < .05. Regarding the Bayesian paradigm, the Bayes Factor (BF) indicates the ratio between the model testing the alternative against the null hypothesis. Customarily, BFs>∼3 provide evidence in favor of the alternative hypothesis, while BFs <∼0.3 indicate evidence in favor of the null hypothesis (van Doorn et al., 2021).

Pre to post-variation rates were tested against 0 using one sample t-test and compared between sessions using paired t-tests. We performed repeated measures ANOVAs to assess the fatigue effect in both the EMG and EEG data, considering SESSION (S10, S12) and BIN (1 to 10) as fixed factors. The same analyses were performed on the Granger results but with a BIN factor reduced to 5 bins. ANOVAs’ results are presented with partial eta squared (η2) and t-test with Cohen’s d (Lakens, 2013). Correlations were performed using Pearson’s r.

All the statistical analyses were performed in JASP (2023, version 0.18.2) (Love et al., 2019), considering default uniform prior distributions for the Bayesian framework.

## V. Acknowledgments

We thank Maxime Picquet, Romain Martinie, Khiara Vorzillo, and Nina Bertho for their valuable help with the data acquisition. We are also grateful to Laurent Gonthier for his technical support in setting up the online force acquisition card and Nathalie Vayssiere for her assistance in obtaining the ethical agreement.

## References

Bailey, S. P., Hall, E. E., Folger, S. E., & Miller, P. C. (2008). Changes in EEG during graded exercise on a recumbent cycle ergometer. Journal of Sports Science & Medicine, 7(4), 505–511.

Barnett, L., & Seth, A. K. (2014). The MVGC multivariate Granger causality toolbox : A new approach to Granger-causal inference. Journal of Neuroscience Methods, 223, 50–68. 10.1016/j.jneumeth.2013.10.018

Barone, J., & Rossiter, H. E. (2021). Understanding the Role of Sensorimotor Beta Oscillations. Frontiers in Systems Neuroscience, 15, 655886. 10.3389/fnsys.2021.655886

Benwell, N. M., Mastaglia, F. L., & Thickbroom, G. W. (2007). Changes in the functional MR signal in motor and non-motor areas during intermittent fatiguing hand exercise. Experimental Brain Research, 182(1), 93–97. 10.1007/s00221-007-0973-5

Bigland-Ritchie, B. (1981). EMG/force relations and fatigue of human voluntary contractions. Exercise and Sport Sciences Reviews, 9, 75–117.

Bigliassi, M. (2015). Corollary discharges and fatigue-related symptoms : The role of attentional focus. Frontiers in Psychology, 6, 1002. 10.3389/fpsyg.2015.01002

Bigliassi, M., & Filho, E. (2022). Functional significance of the dorsolateral prefrontal cortex during exhaustive exercise. Biological Psychology, 175, 108442. 10.1016/j.biopsycho.2022.108442

Buschman, T. J., & Miller, E. K. (2007). Top-down versus bottom-up control of attention in the prefrontal and posterior parietal cortices. *Science (New York*, N.Y.), 315(5820), 1860–1862. 10.1126/science.1138071

Camparo, S., Maymin, P., Park, C., Yoon, S., Zhang, C., Lee, Y., & Langer, E. (2022). The fatigue illusion : The physical effects of mindlessness. Humanities and Social Sciences Communications, 9. 10.1057/s41599-022-01323-0

Ciria, L. F., Perakakis, P., Luque-Casado, A., & Sanabria, D. (2018). Physical exercise increases overall brain oscillatory activity but does not influence inhibitory control in young adults. NeuroImage, 181, 203–210. 10.1016/j.neuroimage.2018.07.009

Crabbe, J. B., & Dishman, R. K. (2004). Brain electrocortical activity during and after exercise : A quantitative synthesis. Psychophysiology, 41(4), 563–574. 10.1111/j.1469-8986.2004.00176.x

Dal Maso, F., Longcamp, M., & Amarantini, D. (2012). Training-related decrease in antagonist muscles activation is associated with increased motor cortex activation : Evidence of central mechanisms for control of antagonist muscles. Experimental Brain Research, 220(3–4), 287–295. 10.1007/s00221-012-3137-1

Delorme, A., & Makeig, S. (2004). EEGLAB : An open source toolbox for analysis of single-trial EEG dynamics including independent component analysis. Journal of Neuroscience Methods, 134(1), 9–21. 10.1016/j.jneumeth.2003.10.009

Denis, G. (2020). Implication des dimensions neurocognitives dans le maintien de l’effort physique au travers du rôle endossé par le cortex préfrontal et de la perspective coûts/bénéfices. Université Côte d’Azur.

Enders, H., Cortese, F., Maurer, C., Baltich, J., Protzner, A. B., & Nigg, B. M. (2016). Changes in cortical activity measured with EEG during a high-intensity cycling exercise. Journal of Neurophysiology, 115(1), 379–388. 10.1152/jn.00497.2015

Engel, A. K., & Fries, P. (2010). Beta-band oscillations—Signalling the status quo? Current Opinion in Neurobiology, 20(2), 156–165. 10.1016/j.conb.2010.02.015

Enoka, R. M., & Stuart, D. G. (1992). Neurobiology of muscle fatigue. Journal of Applied Physiology (Bethesda, Md.: 1985), 72(5), 1631–1648. 10.1152/jappl.1992.72.5.1631

Formaggio, E., Storti, S. F., Avesani, M., Cerini, R., Milanese, F., Gasparini, A., Acler, M., Pozzi Mucelli, R., Fiaschi, A., & Manganotti, P. (2008). EEG and FMRI coregistration to investigate the cortical oscillatory activities during finger movement. Brain Topography, 21(2), 100–111. 10.1007/s10548-008-0058-1

Forrester, L. W., Hanley, D. F., & Macko, R. F. (2006). Effects of treadmill exercise on transcranial magnetic stimulation-induced excitability to quadriceps after stroke. Archives of Physical Medicine and Rehabilitation, 87(2), 229–234. 10.1016/j.apmr.2005.10.016

Gandevia, S. C. (2001). Spinal and supraspinal factors in human muscle fatigue. Physiological Reviews, 81(4), 1725–1789. 10.1152/physrev.2001.81.4.1725

Hosang, L., Mouchlianitis, E., Guérin, S., & Karageorghis, C. (2022). Effects of exercise on electroencephalography-recorded neural oscillations : A systematic review. International Review of Sport and Exercise Psychology, 1–54. 10.1080/1750984X.2022.2103841

Izhikevich, E. M. (2003). Simple model of spiking neurons. IEEE Transactions on Neural Networks, 14(6), 1569–1572. 10.1109/TNN.2003.820440

Izhikevich, E. M., Gally, J. A., & Edelman, G. M. (2004). Spike-timing dynamics of neuronal groups. Cerebral Cortex (New York, N.Y.: 1991), 14(8), 933–944. 10.1093/cercor/bhh053

Jenkinson, N., & Brown, P. (2011). New insights into the relationship between dopamine, beta oscillations and motor function. Trends in Neurosciences, 34(12), 611–618. 10.1016/j.tins.2011.09.003

Jiang, Z., Wang, X.-F., Kisiel-Sajewicz, K., Yan, J. H., & Yue, G. H. (2012). Strengthened functional connectivity in the brain during muscle fatigue. NeuroImage, 60(1), 728–737. 10.1016/j.neuroimage.2011.12.013

Joundi, R. A., Jenkinson, N., Brittain, J.-S., Aziz, T. Z., & Brown, P. (2012). Driving oscillatory activity in the human cortex enhances motor performance. Current Biology: CB, 22(5), 403–407. 10.1016/j.cub.2012.01.024

Kilavik, B. E., Zaepffel, M., Brovelli, A., MacKay, W. A., & Riehle, A. (2013). The ups and downs of β oscillations in sensorimotor cortex. Experimental Neurology, 245, 15–26. 10.1016/j.expneurol.2012.09.014

Kim, G., Ahad, M. A., Ferdjallah, M., & Harris, G. F. (2007). Correlation of muscle fatigue indices between intramuscular and surface EMG signals. Proceedings 2007 IEEE SoutheastCon, 378–382. 10.1109/SECON.2007.342928

Kubitz, K. A., & Mott, A. A. (1996). EEG power spectral densities during and after cycle ergometer exercise. Research Quarterly for Exercise and Sport, 67(1), 91–96. 10.1080/02701367.1996.10607929

Lakens, D. (2013). Calculating and reporting effect sizes to facilitate cumulative science : A practical primer for t-tests and ANOVAs. Frontiers in Psychology, 4, 863. 10.3389/fpsyg.2013.00863

Lebesque, L., Scaglioni, G., & Martin, A. (2022). The impact of submaximal fatiguing exercises on the ability to generate and sustain the maximal voluntary contraction. Frontiers in Physiology, 13, 970917. 10.3389/fphys.2022.970917

Lin, M.-A., Meng, L.-F., Ouyang, Y., Chan, H.-L., Chang, Y.-J., Chen, S.-W., & Liaw, J.-W. (2021). Resistance-induced brain activity changes during cycle ergometer exercises. *BMC Sports Science*, Medicine & Rehabilitation, 13(1), 27. 10.1186/s13102-021-00252-w

Little, S., & Brown, P. (2014). The functional role of beta oscillations in Parkinson’s disease. Parkinsonism & Related Disorders, 20 *Suppl 1*, S44–48. 10.1016/S1353-8020(13)70013-0

Liu, J. Z., Lewandowski, B., Karakasis, C., Yao, B., Siemionow, V., Sahgal, V., & Yue, G. H. (2007). Shifting of activation center in the brain during muscle fatigue : An explanation of minimal central fatigue? NeuroImage, 35(1), 299–307. 10.1016/j.neuroimage.2006.09.050

Liu, J. Z., Shan, Z. Y., Zhang, L. D., Sahgal, V., Brown, R. W., & Yue, G. H. (2003). Human brain activation during sustained and intermittent submaximal fatigue muscle contractions : An FMRI study. Journal of Neurophysiology, 90(1), 300–312. 10.1152/jn.00821.2002

Love, J., Selker, R., Marsman, M., Jamil, T., Dropmann, D., Verhagen, A. J., Ly, A., Gronau, Q., Šmíra, M., Epskamp, S., Matzke, D., Wild, A., Knight, P., Rouder, J., Morey, R., & Wagenmakers, E.-J. (2019). JASP : Graphical Statistical Software for Common Statistical Designs. Journal of Statistical Software, 88. 10.18637/jss.v088.i02

Lundqvist, M., Miller, E. K., Nordmark, J., Liljefors, J., & Herman, P. (2024). Beta : Bursts of cognition. Trends in Cognitive Sciences, S1364-6613(24)00077–9. 10.1016/j.tics.2024.03.010

Masson, M. E. J. (2011). A tutorial on a practical Bayesian alternative to null-hypothesis significance testing. Behavior Research Methods, 43(3), 679–690. 10.3758/s13428-010-0049-5

Matta, P.-M., Glories, D., Alamia, A., Baurès, R., & Duclay, J. (2024). Mind over muscle? Time manipulation improves physical performance by slowing down the neuromuscular fatigue accumulation. Psychophysiology, 61(4), e14487. 10.1111/psyp.14487

McMorris, T., Barwood, M., & Corbett, J. (2018). Central fatigue theory and endurance exercise : Toward an interoceptive model. Neuroscience and Biobehavioral Reviews, 93, 93–107. 10.1016/j.neubiorev.2018.03.024

Merletti, R., Lo Conte, L. R., & Orizio, C. (1991). Indices of muscle fatigue. Journal of Electromyography and Kinesiology: Official Journal of the International Society of Electrophysiological Kinesiology, 1(1), 20–33. 10.1016/1050-6411(91)90023-X

Miller, E. K., & Cohen, J. D. (2001). An integrative theory of prefrontal cortex function. Annual Review of Neuroscience, 24, 167–202. 10.1146/annurev.neuro.24.1.167

Neuper, C., & Pfurtscheller, G. (2001a). Event-related dynamics of cortical rhythms : Frequency-specific features and functional correlates. International Journal of Psychophysiology: Official Journal of the International Organization of Psychophysiology, 43(1), 41–58. 10.1016/s0167-8760(01)00178-7

Neuper, C., & Pfurtscheller, G. (2001b). Evidence for distinct beta resonance frequencies in human EEG related to specific sensorimotor cortical areas. Clinical Neurophysiology: Official Journal of the International Federation of Clinical Neurophysiology, 112(11), 2084–2097. 10.1016/s1388-2457(01)00661-7

Okamoto, M., Dan, H., Sakamoto, K., Takeo, K., Shimizu, K., Kohno, S., Oda, I., Isobe, S., Suzuki, T., Kohyama, K., & Dan, I. (2004). Three-dimensional probabilistic anatomical cranio-cerebral correlation via the international 10-20 system oriented for transcranial functional brain mapping. NeuroImage, 21(1), 99–111. 10.1016/j.neuroimage.2003.08.026

Oswal, A., Brown, P., & Litvak, V. (2013). Synchronized neural oscillations and the pathophysiology of Parkinson’s disease. Current Opinion in Neurology, 26(6), 662–670. 10.1097/WCO.0000000000000034

Pfurtscheller, G., & Lopes da Silva, F. H. (1999). Event-related EEG/MEG synchronization and desynchronization : Basic principles. Clinical Neurophysiology: Official Journal of the International Federation of Clinical Neurophysiology, 110(11), 1842–1857. 10.1016/s1388-2457(99)00141-8

Pfurtscheller, G., Neuper, C., Pichler-Zalaudek, K., Edlinger, G., & Lopes da Silva, F. H. (2000). Do brain oscillations of different frequencies indicate interaction between cortical areas in humans? Neuroscience Letters, 286(1), 66–68. 10.1016/s0304-3940(00)01055-7

Pfurtscheller, G., Stancák, A., & Neuper, C. (1996). Post-movement beta synchronization. A correlate of an idling motor area? Electroencephalography and Clinical Neurophysiology, 98(4), 281–293. 10.1016/0013-4694(95)00258-8

Pion-Tonachini, L., Kreutz-Delgado, K., & Makeig, S. (2019). ICLabel : An automated electroencephalographic independent component classifier, dataset, and website. NeuroImage, 198, 181–197. 10.1016/j.neuroimage.2019.05.026

Pogosyan, A., Gaynor, L. D., Eusebio, A., & Brown, P. (2009). Boosting cortical activity at Beta-band frequencies slows movement in humans. Current Biology: CB, 19(19), 1637–1641. 10.1016/j.cub.2009.07.074

Poulet, J. F. A., & Hedwig, B. (2006). The cellular basis of a corollary discharge. *Science (New York*, N.Y.), 311(5760), 518–522. 10.1126/science.1120847

Robertson, C. V., & Marino, F. E. (2015). Prefrontal and motor cortex EEG responses and their relationship to ventilatory thresholds during exhaustive incremental exercise. European Journal of Applied Physiology, 115(9), 1939–1948. 10.1007/s00421-015-3177-x

Robertson, C. V., & Marino, F. E. (2016). A role for the prefrontal cortex in exercise tolerance and termination. Journal of Applied Physiology (Bethesda, Md.: 1985), 120(4), 464–466. 10.1152/japplphysiol.00363.2015

Rönnefarth, M., Jooß, A., Haberbosch, L., Köhn, A., Fleischmann, R., Brandt, S., & Schmidt, S. (2017). Frequency specific modulation of motor fatigue by beta- and gamma-tACS. Clinical Neurophysiology, 128(10), e356. 10.1016/j.clinph.2017.06.133

Rossiter, H. E., Davis, E. M., Clark, E. V., Boudrias, M.-H., & Ward, N. S. (2014). Beta oscillations reflect changes in motor cortex inhibition in healthy ageing. NeuroImage, 91(100), 360–365. 10.1016/j.neuroimage.2014.01.012

Sallard, E., Tallet, J., Thut, G., Deiber, M.-P., & Barral, J. (2014). Post-switching beta synchronization reveals concomitant sensory reafferences and active inhibition processes. Behavioural Brain Research, 271, 365–373. 10.1016/j.bbr.2014.05.070

Schmidt, R., Herrojo Ruiz, M., Kilavik, B. E., Lundqvist, M., Starr, P. A., & Aron, A. R. (2019). Beta Oscillations in Working Memory, Executive Control of Movement and Thought, and Sensorimotor Function. The Journal of Neuroscience: The Official Journal of the Society for Neuroscience, 39(42), 8231–8238. 10.1523/JNEUROSCI.1163-19.2019

Seth, A. K., Barrett, A. B., & Barnett, L. (2015). Granger causality analysis in neuroscience and neuroimaging. The Journal of Neuroscience: The Official Journal of the Society for Neuroscience, 35(8), 3293–3297. 10.1523/JNEUROSCI.4399-14.2015

Solis-Escalante, T., Müller-Putz, G. R., Pfurtscheller, G., & Neuper, C. (2012). Cue-induced beta rebound during withholding of overt and covert foot movement. Clinical Neurophysiology: Official Journal of the International Federation of Clinical Neurophysiology, 123(6), 1182–1190. 10.1016/j.clinph.2012.01.013

Stancák, A., Riml, A., & Pfurtscheller, G. (1997). The effects of external load on movement-related changes of the sensorimotor EEG rhythms. Electroencephalography and Clinical Neurophysiology, 102(6), 495–504. 10.1016/s0013-4694(96)96623-0

Stevenson, C. M., Brookes, M. J., & Morris, P. G. (2011). β-Band correlates of the fMRI BOLD response. Human Brain Mapping, 32(2), 182–197. 10.1002/hbm.21016

Stoll, F. M., Wilson, C. R. E., Faraut, M. C. M., Vezoli, J., Knoblauch, K., & Procyk, E. (2016). The Effects of Cognitive Control and Time on Frontal Beta Oscillations. *Cerebral Cortex (New York*, N.Y.: 1991), *26*(4), 1715–1732. 10.1093/cercor/bhv006

Suviseshamuthu, E., Handiru, V., Allexandre, D., Hoxha, A., Saleh, S., & Yue, G. (2021). EEG-Based Spectral Analysis Showing Brainwave Changes Related to Modulating Progressive Fatigue During a Prolonged Intermittent Motor Task. 10.1101/2021.09.08.458591

Tanaka, M., & Watanabe, Y. (2012). Supraspinal regulation of physical fatigue. Neuroscience and Biobehavioral Reviews, 36(1), 727–734. 10.1016/j.neubiorev.2011.10.004

Taylor, J. L., Amann, M., Duchateau, J., Meeusen, R., & Rice, C. L. (2016). Neural Contributions to Muscle Fatigue : From the Brain to the Muscle and Back Again. Medicine and Science in Sports and Exercise, 48(11), 2294–2306. 10.1249/MSS.0000000000000923

van Doorn, J., van den Bergh, D., Böhm, U., Dablander, F., Derks, K., Draws, T., Etz, A., Evans, N. J., Gronau, Q. F., Haaf, J. M., Hinne, M., Kucharský, Š., Ly, A., Marsman, M., Matzke, D., Gupta, A. R. K. N., Sarafoglou, A., Stefan, A., Voelkel, J. G., & Wagenmakers, E.-J. (2021). The JASP guidelines for conducting and reporting a Bayesian analysis. Psychonomic Bulletin & Review, 28(3), 813–826. 10.3758/s13423-020-01798-5

Wach, C., Krause, V., Moliadze, V., Paulus, W., Schnitzler, A., & Pollok, B. (2013). Effects of 10 Hz and 20 Hz transcranial alternating current stimulation (tACS) on motor functions and motor cortical excitability. Behavioural Brain Research, 241, 1–6. 10.1016/j.bbr.2012.11.038

Yang, Q., Fang, Y., Sun, C.-K., Siemionow, V., Ranganathan, V. K., Khoshknabi, D., Davis, M. P., Walsh, D., Sahgal, V., & Yue, G. H. (2009). Weakening of functional corticomuscular coupling during muscle fatigue. Brain Research, 1250, 101–112. 10.1016/j.brainres.2008.10.074

Yuan, H., Liu, T., Szarkowski, R., Rios, C., Ashe, J., & He, B. (2010). Negative covariation between task-related responses in alpha/beta-band activity and BOLD in human sensorimotor cortex : An EEG and fMRI study of motor imagery and movements. NeuroImage, 49(3), 2596–2606. 10.1016/j.neuroimage.2009.10.028

